# Alexidine dihydrochloride has broad spectrum activities against diverse fungal pathogens

**DOI:** 10.1101/429944

**Authors:** Zeinab Mamouei, Abdullah Alqarihi, Shakti Singh, Shuying Xu, Michael K. Mansour, Ashraf S Ibrahim, Priya Uppuluri

## Abstract

Invasive fungal infections due to *Candida albicans*, *Aspergillus fumigatus* and *Cryptococcus neoformans*, constitute a substantial threat to hospitalized, immunocompromised patients. Further, the presence of drug-recalcitrant biofilms on medical devices, and emergence of drug-resistant fungi such as *Candida auris*, introduce treatment challenges with current antifungal drugs. Worse, currently there is no approved drug capable of obviating preformed biofilms which increases the chance of infection relapses. Here, we screened a small molecule Prestwick Chemical Library, consisting of 1200 FDA approved off-patent drugs, against *C. albicans*, *C. auris* and *A. fumigatus*, to identify those that inhibit growth of all three pathogens. Inhibitors were further prioritized for their potency against other fungal pathogens, and their ability to kill preformed biofilms. Our studies identified the bis-biguanide Alexidine dihydrochloride (AXD), as a drug with the highest antifungal and anti-biofilm activity against a diverse range of fungal pathogens. Finally, AXD significantly potentiated the efficacy of fluconazole against biofilms, displayed low mammalian cell toxicity, and eradicated biofilms growing in mice central venous catheters *in vivo*, highlighting its potential as a pan-antifungal drug.

**Importance:** The prevalence of fungal infections has seen a rise in the past decades due to advances in modern medicine leading to an expanding population of device-associated and immunocompromised patients. Furthermore, the spectrum of pathogenic fungi has changed, with the emergence of multi-drug resistant strains such as *C. auris*. High mortality related to fungal infections point to major limitations of current antifungal therapy, and an unmet need for new antifungal drugs. We screened a library of repurposed FDA approved inhibitors to identify compounds with activities against a diverse range of fungi, in varied phases of growth. The assays identified Alexidine dihydrochloride (AXD) to have pronounced antifungal activity including against preformed biofilms, at concentrations lower than mammalian cell toxicity. AXD potentiated the activity of fluconazole and amphotericin B against *Candida* biofilms *in vitro*, and prevented biofilm growth *in vivo*. Thus AXD has the potential to be developed as a pan-antifungal, anti-biofilm drug.

## Introduction

Fungal pathogens responsible for invasive fungal infections (IFIs) are a leading cause of human mortality, killing approximately one and a half million people every year, despite treatment with antifungal drugs (1). Of concern, the current incidence of fungal-related deaths is reported to be even higher than mortality due to tuberculosis or malaria (2). A vast majority of IFIs result from species belonging to *Cryptococcus*, *Candida* or *Aspergillus* (3). However, fungi such as molds other than *Aspergillus*, and non-*albicans Candida* species including the multi-drug resistant pathogen *C. auris*, are becoming increasingly frequent and difficult to treat (4). Furthermore, other IFIs such as those due to Mucorales cause highly angioinvasive and tissue-destructive infections which in many cases have mortality rates close to 100% (2).

The challenge in treatment of IFIs is directly linked to an ever expanding population of immunocompromised patients requiring modern medical interventions, and a paucity of currently approved antifungal agents (5, 6). Indwelling medical devices infected with fungi, develop biofilms that are notoriously resistant to all classes of antifungal drugs, and serve as a reservoir of infectious cells with direct access to the vasculature (7, 8). Current therapeutic armamentarium for IFIs is sparse, including only three classes of antifungal agents: polyenes, azoles, and echinocandins. These drugs have drawbacks including significant limitations in spectrum of activity, human toxicity and emergence of drug resistance, thereby underscoring a need for development of new antifungal agents (9, 10).

To fulfil this unmet need, we employed a high throughput screening assay (HTS), to screen and characterize FDA (U.S. Food and Drug Administration)-approved, off patent library drugs for their abilities to kill/inhibit three of the most invasive and drug-resistant human pathogenic fungi, *C. albicans*, *C. auris* and *A. fumigatus*. This assay allowed us to identify core fungicidal molecules against all the three pathogens. One of the leading compounds identified was a bis-biguanide dihydrochloride called alexidine dihydrochloride (AXD). AXD is an anti-cancer drug that targets a mitochondrial tyrosine phosphatase PTPMT1 in mammalian cells, and causes mitochondrial apoptosis (11). We found that AXD not only inhibited planktonic growth, but also prevented biofilm formation, as well as killed biofilms formed by a variety of drug resistant and susceptible isolates of diverse fungal organisms. Further, when used in combination, AXD reduced the MIC of fluconazole and amphotericin B, and rendered them efficacious against drug resistant *C. albicans* biofilms. Finally, anti-biofilm property of AXD was also recapitulated in an *in vivo* mouse central venous catheter model of *C. albicans* biofilm formation. Overall, our studies warrant the further development of AXD as a pan-fungal anti-biofilm drug, which could be used in combination therapeutics against diverse fungal pathogens.

## Results and discussion

### High throughput screening (HTS) for identification of antifungal molecules

We used a HTS assay to test the ability of a commercially available, small molecule library containing 1233 FDA approved compounds (New Prestwick Chemical [NPW] Library). We reckoned that repositioning existing off-patent drugs with known human safety and bioavailability profiles, can accelerate the antifungal drug-discovery process without undergoing the arduous FDA approval process. These compounds were screened to identify a core set of inhibitors and fungicidals against *C. albicans*, *A. fumigatus* and *C. auris*. The former two fungi represent two of the top four fungal pathogens causing IFIs with 40-70% mortality rates (3). *C. auris* is a newly emerging fungus that represents a serious global health threat due to its multi-drug resistant properties (12). We used cell viability as a parameter for prioritizing the broad spectrum FDA-approved molecules as lead drugs for developing pan-fungal therapeutics. The purpose was to first identify a core set of molecules that could inhibit a diverse collection of fungi spanning different genus and species, under planktonic growth conditions.

HTS was performed in a 384-well plate screening format, where the NPW library was screened against planktonic yeast or spore suspensions of the three fungal organisms, at a single concentration of 10 μM. The spectrum of activity of these drugs was compared to clinically used azole drugs (fluconazole or voriconazole) at a concentration ranging from 0.03 to 32 μg/ml. MIC of drugs were determined in agreement with the CLSI M27-A3 (for yeast) and M38-A2 (for filamentous fungi) reference standards for antifungal susceptibility testing (13, 14). After three days of incubation at 37oC, turbidity of the wells (OD600) was measured and molecules displaying >50% reduction in turbidity compared to control non-drug treated wells (MIC50) were considered as primary “hits”. Z′ factor, was calculated as a parameter of HTS screening quality, and an average Z′ factor of 0.75 was computed for our assays (a value of >0.5 represents an excellent quality of HTS)(15).

From this hit-list, a core set of molecules that inhibited planktonic growth of all three fungi as identified by >50% growth inhibition measured by MIC were identified and shortlisted. *C. albicans* was sensitive to fluconazole at concentration <0.125-0.25 μg/ml, as has been reported previously (16), while consistent with its drug resistant nature, *C. auris* was resistant to fluconazole with MIC >16 μg/ml (Table 2). *A. fumigatus* succumbed to voriconazole at 0.25 μg/ml, similar to previously reported anti-fungal drug susceptibility studies (16). Recently, Siles *et al.* investigated the ability of NPW to specifically inhibit *C. albicans* biofilms, and revealed 38 pharmacologically active agents against the fungus (17). While our study also identified a number of molecules individually inhibiting the three fungi, respectively (Table. 1), only the following six compounds were successful at inhibiting all three organisms: chloroxin, thimerosal, alexidine dihydrochloride, haloprogin, clioquinol and butenafine hydrochloride (Table 1). NPW contains a number of antifungal drugs, such as imidazoles’, triazoles and polyene class of drugs. *C. auris* was by far the most resistant fungus, inert against the azoles and polyenes in the library. The six molecules were further evaluated for their ability to curtail biofilm formation as well as kill pre-formed biofilms developed by the three fungi.

**Table 1:** Table 1. Hits obtained from primary screening of the New Prestwick Chemical Library against planktonic cells of *C. albicans*, *A.fumigatus* and *C. auris*

| **<u>C. albicans</u>** | **<u>A. fumigatus</u>** |
| --- | --- |
| Alexidine dihydrochloride | Alexidine dihydrochloride |
| Amphotericin B | Butenafine Hydrochloride |
| Antimycin A | Isoxsuprine hydrochloride |
| Butenafine Hydrochloride | Clioquinol |
| Chloroxine | Thimerosal |
| Ciclopirox ethanolamine | Dequalinium dichloride |
| Clioquinol | Pyrvinium pamoate |
| Clotrimazole | Haloprogin |
| Dequalinium dichloride | Chlorhexidine |
| Econazole nitrate | Voriconazole |
| Enilconazole | Amphotericin B |
| Fluconazole | Antimycin A |
| Flucytosine | Econazole nitrate |
| Haloprogin | Enilconazole |
| Isoconazole | Methyl benzethonium chloride |
| Isoxsuprine hydrochloride | Tioconazole |
| Itraconazole | **<u>C. auris</u>** |
| Ketoconazole | Alexidine dihydrochloride |
| Methyl benzethonium chloride | Butenafine Hydrochloride |
| Pyrvinium pamoate | Chloroxine |
| Sertaconazole nitrate | Clioquinol |
| Sulconazole nitrate | Thimerosal |
| Terconazole | Haloprogin |
| Thimerosal |  |
| Thonzonium bromide |  |
| Tioconazole |  |
| Voriconazole |  |

**Table 2.**
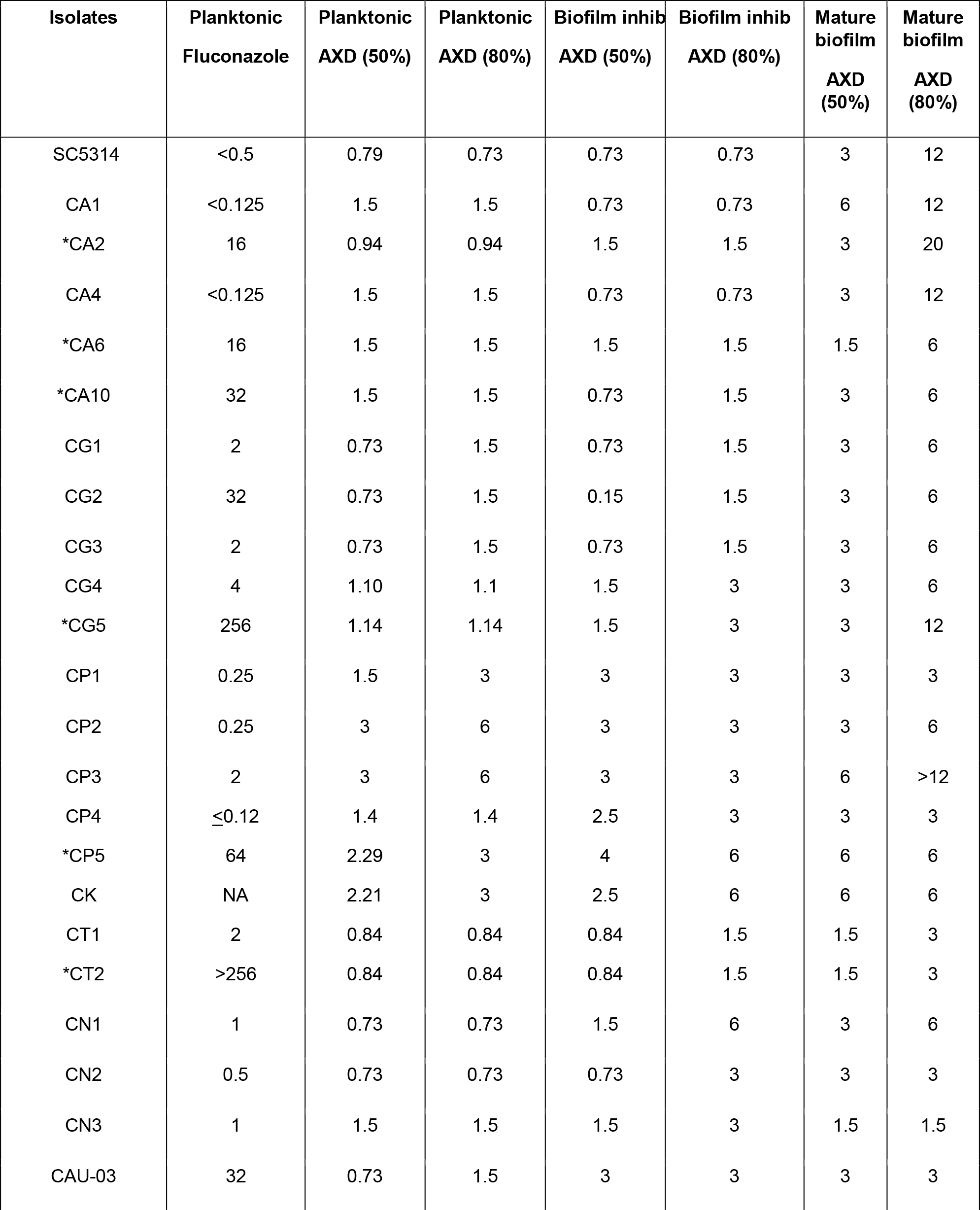
MIC of AXD against clinical isolates of different fungal species, *vs* fluconazole or voriconazole. Values are in μg/ml. CA=*C. albicans*, CG=*C. glabrata,* CP=*C. parapsilosis,* CK=*C. krusei,* CN=*C. neoformans,* AF=*A. fumigatus*

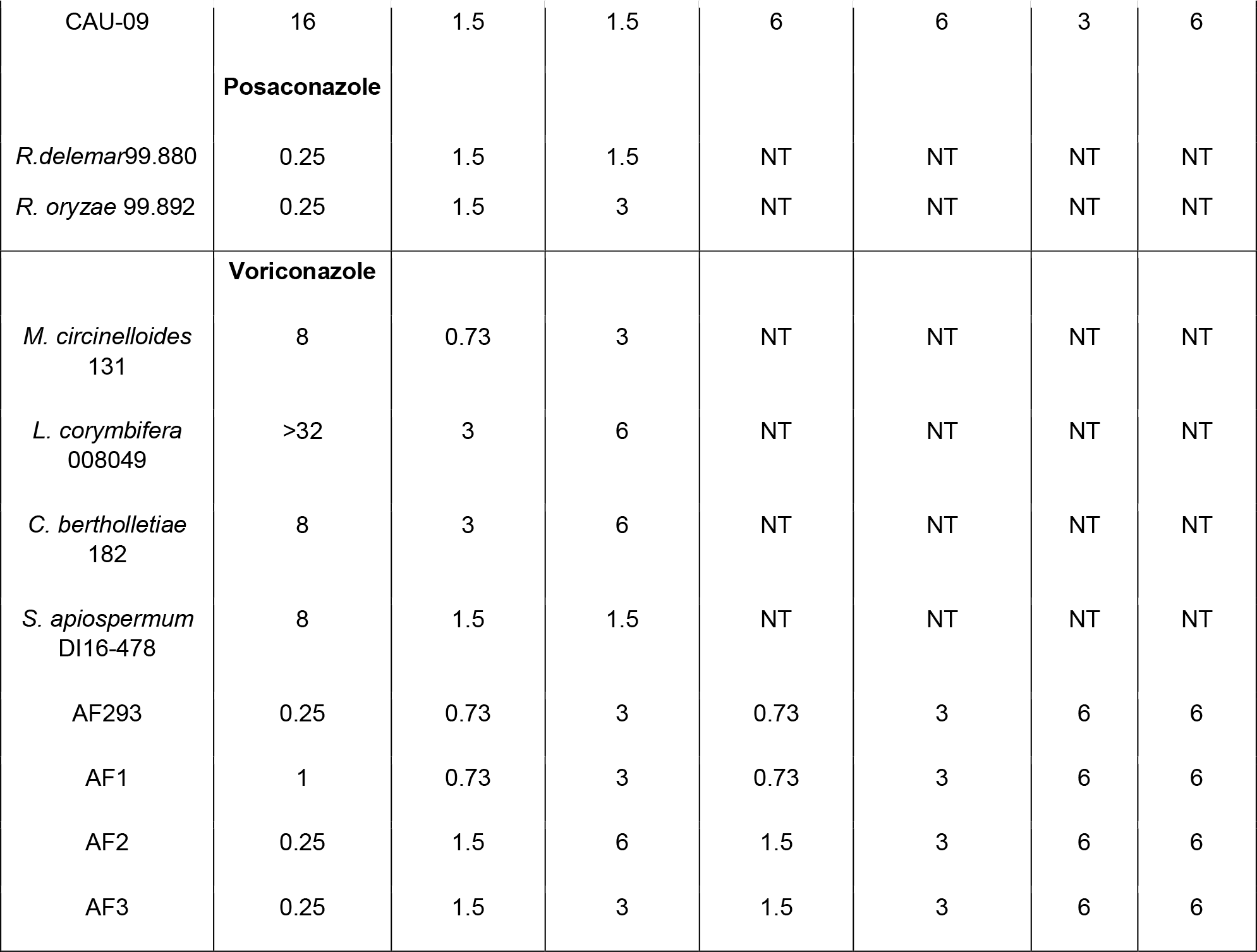

### Secondary assays for determination of anti-biofilm activity

Wells with *C. albicans*, *C. auris* and *A. fumigatus* were either treated with inhibitors at the time of yeast/spore inoculation (start of biofilm initiation), or allowed to grow without drugs for 48 h to allow biofilm development (mature biofilm). This assay was performed in a 96-well microtiter plate assay, as previously reported by us (18). For the effect on formation of biofilm, all six inhibitors could inhibit biofilm formation in the three fungal organisms, as adjudged by a significant decrease in turbidity of the media in the wells 48 h following incubation with the drugs (data not shown). However, only two drugs, alexidine dihydrochloride and thimerosal could significantly kill 80% of mature biofilm community at the tested concentration of <10 μM (Fig 1A and **S1A**). We chose to focus our attention to studying alexidine dihydrochloride (AXD), since it was more attractive with respect to drug development, having indications for use as an antibacterial and antiplaque agent and with limited side effects (19–21).

**Figure 1.**
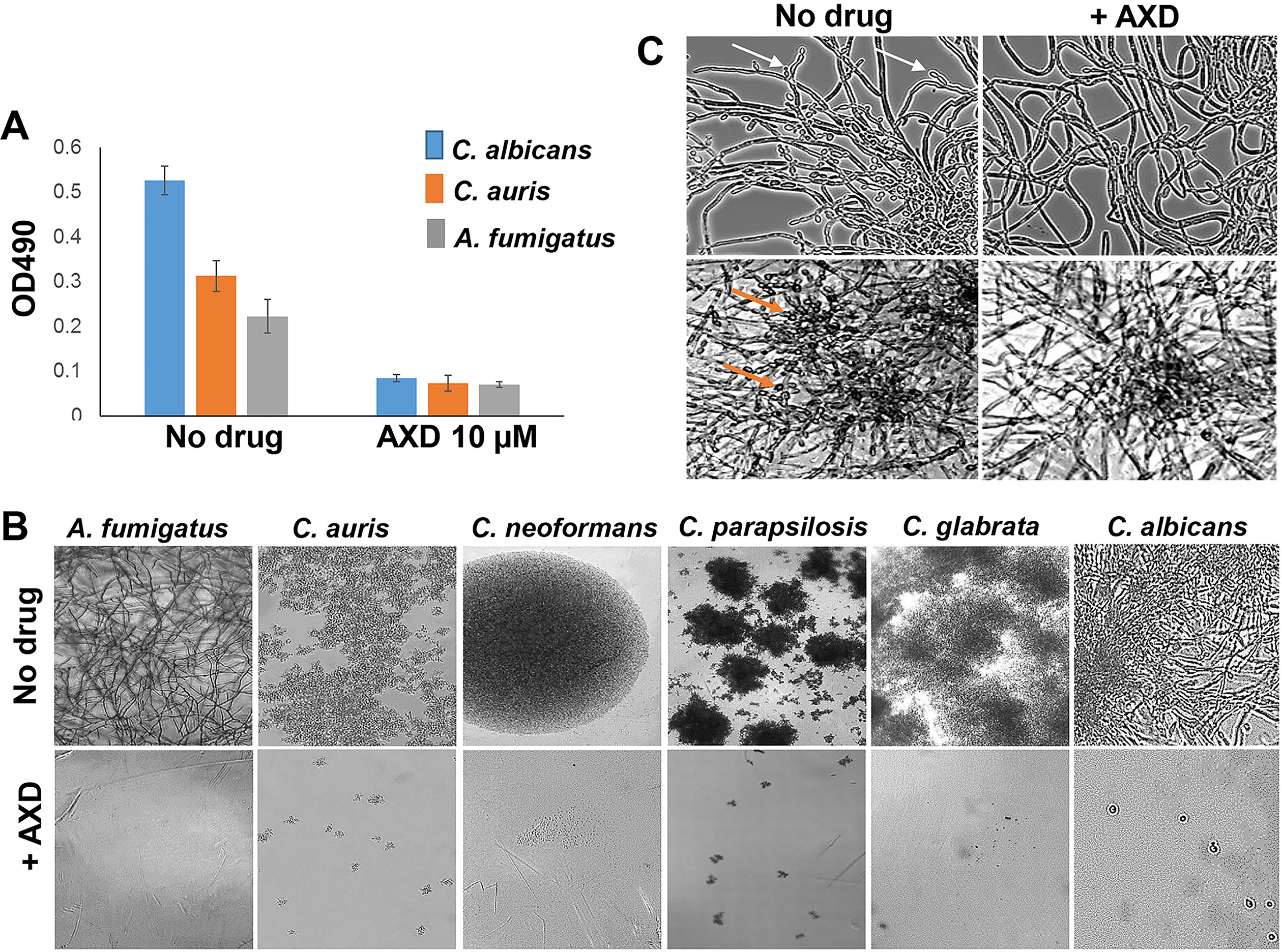
Inhibition of biofilm growth, *C. albicans* biofilm dispersal, and abrogation of planktonic growth in diverse fungi by alexidine dihydrochloride (AXD): A) Fungal cells were allowed to form a biofilm for 48 h and treated for 24 h with 10 μM AXD. Biofilm inhibition was as determined by XTT reading (OD490). B) Fungal yeast cells or spores were incubated under different concentrations of AXD under planktonic conditions. Inhibition of growth and filamentation of the fungi visualized by phase contrast microscopy (20X magnification), at their respective AXD MIC80 concentrations. C) *C. albicans* planktonic hyphae (top two panels) and biofilms (bottom two panels) were treated for 12 h with 150 ng/ml of AXD. AXD inhibited lateral yeast production from hyphal cells and hyphal layers of biofilms, as visualized microscopically. Arrows point to lateral yeasts.

### Dose response assays of AXD

AXD was first evaluated in planktonic and biofilm dose response assays, and further tested for its ability to inhibit growth of other fungal pathogens including drug resistant clinical isolates, a number of non-*albicans Candida* spp, and members of the Mucorales family. The results for this study are described in Table. 2 which lists the fungal strains used for evaluation of the efficacy of AXD, under three different growth conditions – planktonic, biofilm inhibition and preformed biofilms, compared to the MIC of the control azole antifungal drugs. For example, AXD displayed activity against most *Candida* spp; MIC values of ≤1.5 μg/ml were observed for all isolates tested under planktonic conditions, with the exception of *C. parapsilosis* and *C. krusei.* Interestingly, AXD also displayed striking activity against clinically relevant fluconazole-resistant *Candida* isolates: *C. albicans* (CA2, CA6 and CA10), *C. glabrata* (CG2, CG5), *C. parapsilosis* (CP5) and *C. auris* (CAU-09, CAU-03). Furthermore, the MIC values of AXD against C. *neoformans was* comparable to the MIC values for fluconazole.

In case of filamentous fungi, low AXD MIC50 values of 1.5-3 μg/ml was observed for all filamentous fungi (Mucorales and *Aspergillus* spp., plates read at 48 h), including the molds *L. corymbifera* and *S. apiospermum* (read at 72 h) that have poor outcome with current clinically available antifungal drugs (22). Inhibition of planktonic growth by AXD, monitored microscopically, revealed a complete inhibition of filamentation or proliferation of the imaged fungi (**Fig 1B**). Of particular importance was the finding that AXD was able to decimate at low concentrations (1.5-6 μg/ml), mature biofilms of *Candida*, *Cryptococcus* and *Aspergillus* spp. that are known to be resistant to almost all classes of antifungal drugs (Table 2, also see **S2** for AXD activity on *Candida* spp.). In fact, at 10 fold lower concentrations (150 ng/ml) of planktonic MICs’, AXD could inhibit lateral yeast formation and biofilm dispersal in *C. albicans* (**Fig 1C**). Dispersal of lateral yeast cells from a biofilm biomass is the link between contaminated catheters and disseminated candidiasis (8, 23). Inhibition of dispersal with just nano-molar levels of AXD can help seal the biofilm reservoir and curtail further proliferation and robustness of a biofilm.

Alexidine dihydrochloride is a bis(biguanide) in which the common 2-ethylhexyl chain has been attached to each biguanide unit and the two units are linked by a 1,6-hexanediamine chain (**S3A**). This compound initially identified for its anti-bacterial properties, is also found as an inducer of mitochondrial dysfunction and apoptosis (11, 24). AXD has been proposed in several studies as an anti-plaque agent, mouthwash, and with potential to be used in endodontic treatment to eliminate biofilms (19, 20, 25). These reports, along with our present findings, serve as precedents for the development of AXD as an anti-biofilm agent. Another bisbiguanide with structural similarity to AXD, metformin, has recently been shown to have antifungal activity (and synergistic potentiation of clinically used antifungal drugs) against *C. glabrata*, albiet only at physiologically inapt high concentrations (26). AXD, on the other hand is active even at levels as low as 0.75 μg/ml, against an array of fungal species in our current study. In fact, AXD has been reported to have activity against the fungus *C. neoformans*, by targeting phospholipases (27). Whether specific inhibition of fungal phospholipases is the cause of AXD’s antifungal activity against a spectrum of pathogenic fungi, is unknown and remains to be explored in future studies.

### Mammalian cell cytotoxicity assays and synergy of AXD with fluconazole

Considering that AXD displayed enhanced efficacy against fungal organisms, we evaluated the extent of its cell toxicity (CC50) to mammalian cells. Results showed that AXD resulted in 50% killing of HUVECs and lung epithelial cells, at concentrations 5-10 fold higher than the MIC required to kill planktonically growing fungal pathogens (CC50 >7.37 μg/ml vs planktonic MIC50 of 0.73-1.5 μg/ml) (**Fig. 2A, B**). Previous studies have reported similar cytotoxicity levels of AXD against various other cell lines (19, 27, 28). We further tested the toxicity of AXD to a human bone-marrow derived macrophage cell line to understand its effect on the immune cells. AXD displayed a slightly higher toxicity to the macrophages compared to the mammalian tissue cell lines, with a CC50 of over 5 μg/ml (**Fig 2C**). A similar study was also done to test the impact of AXD toxicity on HL60 monocyte proliferation. HL60 cells stained with CFSE were treated with varying concentrations of AXD, or the control PMA (phorbol 12-myristate 13-acetate) as a positive stimulant that induces cellular proliferation. Inhibition of cellular proliferation corresponds to toxicity, and the concentration of AXD that could prevent early cell division in HL60 cells was examined. As expected cells stimulated by PMA showed proliferation, while those treated with the highest dose of AXD (10 μg/ml) did not divide. AXD at 5 μg/ml prevented cell division in the human bone marrow derived HL60 cells (**S3B**). This level of toxicity matched the macrophage CC50 value. We note that concentrations detrimental to host cells are at least 3-4 fold higher than AXD levels required to inhibit planktonic cells of many different fungi, including *C. albicans*. These moderately low cytotoxicity of the FDA approved drug pave the way to a potential repurposing of AXD as an antifungal agent, and warrant its further development into a compound with higher efficacy, bioavailability and less toxicity.

**Figure 2.**
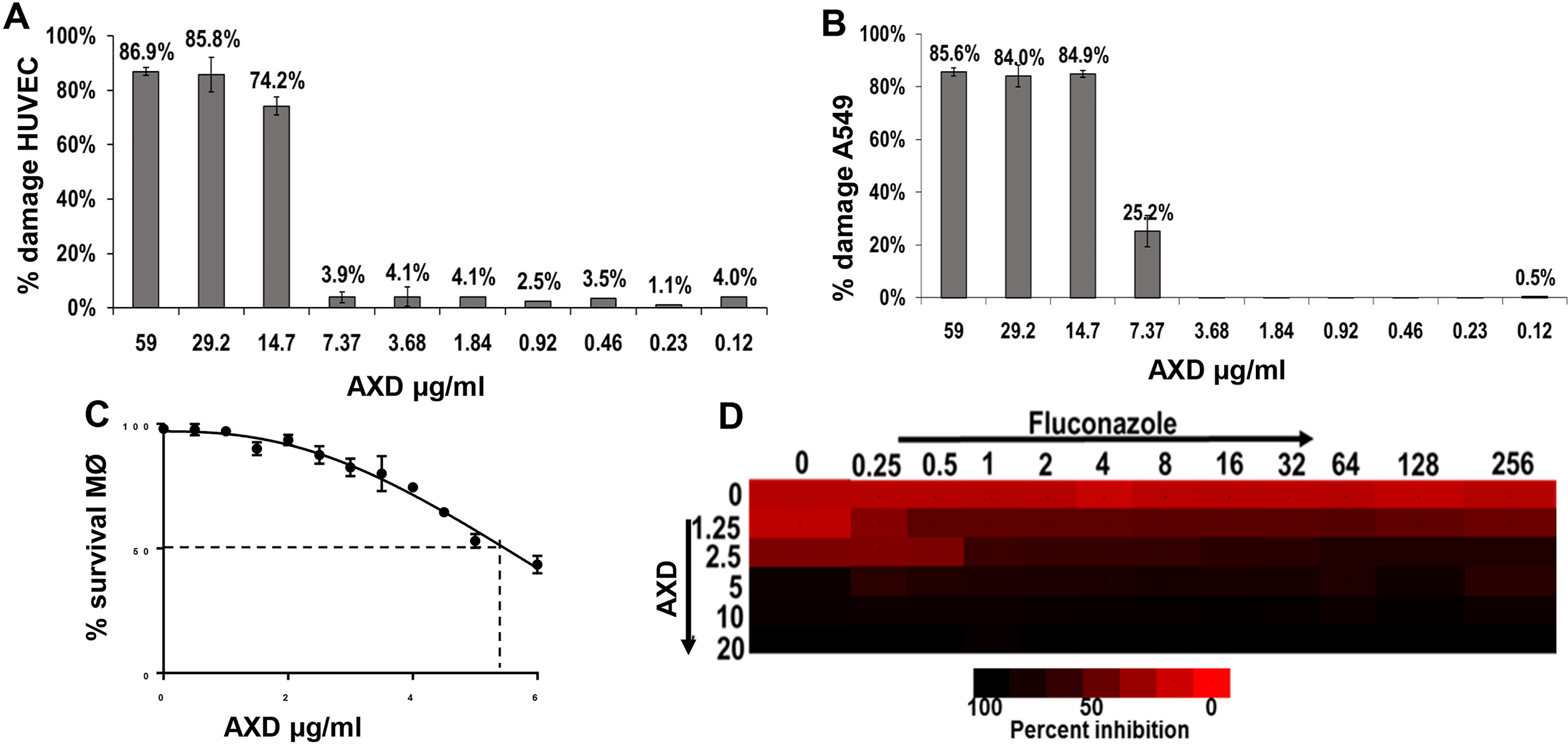
Toxicity of AXD on host cells, and on biofilm killing in combination with fluconazole. Different concentrations of AXD were incubated with HUVEC (A), lung A549 (B), or macrophages (C) for 24 h at 37°C, for testing the CC50 of the drug to the respective cell lines. (D) *C. albicans* biofilms were developed for 48 h and then treated with different concentrations of AXD and fluconazole in a checkerboard format. Metabolic activity of biofilm cells were measured by the XTT assay. Bright red represents growth above the MIC_50_, dull red represents growth at the MIC_50_, and black/dark red represents growth below the MIC_50_.

This inhibitory potential was further highlighted in our studies evaluating synergistic action of AXD in combination with fluconazole, against mature *C. albicans* biofilms. Fluconazole is completely inert against *C. albicans* biofilms, with an MIC50 of >250 μg/ml {this study and (7, 29, 30)}. When used together, AXD at 1.25 μg/ml, strikingly reduced the MIC50 of fluconazole from >256 μg/ml to clinically relevant 1 μg/ml (**Fig 2D**), providing an FIC index of 0.42, that indicated a synergistic interaction (30). These results further emphasize AXD’s prospect as an anti-biofilm agent, especially due to its ability to lower MICs of fluconazole, highlighting the possibility of bringing a biofilm-redundant drug back into clinical use.

### Inhibition of biofilm *in vivo* by AXD

Our studies showed that AXD could arrest growth and kill biofilm cells formed by various *Candida* species, *C. neoformans* and *A. fumigatus* in *in vitro* assays. We next examined the ability of AXD to decimate preformed biofilms in an *in vivo* model. For this study we chose to focus on biofilm formation by *C. albicans*, since a murine biofilm model has been well established in this fungus and used for testing the effects of established and new antifungal agents (31).

The effect of the drugs on the 24 h old biofilms growing in the jugular vein catheters of mice was visualized microscopically, which revealed significantly lower density of the biofilms in catheters treated with AXD and caspofungin, versus the control untreated catheters (**Fig 3A**). In fact, fungal CFU determination revealed that AXD inhibited 67% of fungal biofilm growth and viability, compared to the control untreated biofilms (**Fig 3B**). As expected, caspofungin (an antifungal drug known to be hyperactive against *C. albicans* biofilms) decimated >90% of the biofilm community growing within the catheters. On the other hand, fluconazole (a drug with enhanced activity against planktonic fungi, but with no effectiveness against biofilm cells) was found to reduce biofilms by only 30% (**Fig 3B**). Overall our data shows that AXD can inhibit biofilm growth *in vivo*. A better understanding of the pharmacokinetics/pharmacodynamics of AXD could be invaluable in assessing its utility as a systemic antifungal drug, especially in a disseminated mouse model of fungemia.

**Figure 3.**
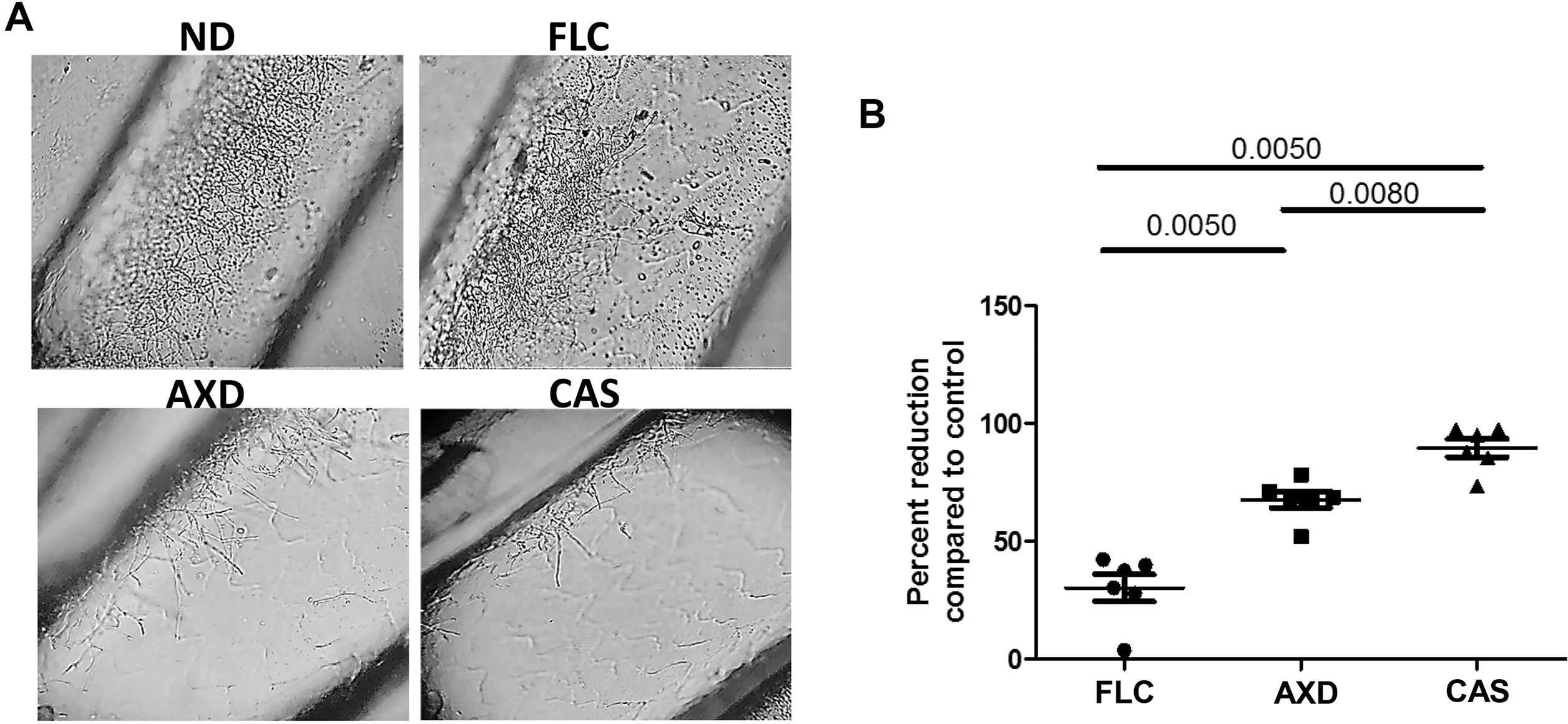
Impact of AXD, fluconazole (FLC) and caspofungin (CAS) as lock therapy against *C. albicans* biofilm cells in an in *vivo* catheter model. (A) Biofilms were grown for 24 h followed by intraluminal drug treatment for 24 h. Following compound exposure, the catheters were removed for microscopy and CFU enumeration. Each of the four panels represent a 40× magnification under phase contrast microscope. Panel columns: no drug treatment (ND): control biofilm treated with saline; FLC, 125-μg/ml fluconazole exposure; AXD, catheters exposed to AXD at 3 μg/ml; CAS, catheters exposed to 0.25 μg/ml caspofungin. (B) Post ND or drug treatment, catheters were cut into pieces, vortexed and sonicated to release adhered cells in sterile PBS and dilutions of the suspension were plated on solid media for CFU enumeration. Results are presented as percent biofilm reduction in drug-treated catheters compared to the untreated catheter-biofilms, and analyzed statistically by using a non-parametric t-test. P value of <0.05 is significant.

In summary, our HTS identified alexidine dihydrochloride to have profound activity against various growth forms of fungi: planktonic, biofilm and biofilm dispersal. AXD was fungicidal to a number of different pathogenic fungi including common as well as emerging drug resistant pathogens. The fact that AXD retains its activity against azole resistant clinical isolates indicates its potential use in recalcitrant fungal infections. Importantly, AXD reduces the MIC of fluconazole-a clinically used first line antifungal drug, ironically considered dispensable for biofilm treatment, thereby pointing to its extended utility as an anti-biofilm combination drug. Perhaps the most intriguing activity of AXD was seen against Mucorales including *Rhizopus*, a species that leads to devastating infections and very poor outcomes in patients, despite conventional antifungal treatment. Furthermore, the drug was also potent against those fungi that are therapeutically unmanageable in clinics with current antifungal agents such as *L. corymbifera* and *S. apiospermum*. Future studies will focus on the mechanism of action of AXD at a molecular level, and evaluate its feasibility as a pan-antifungal drug to combat infections in different clinical settings.

## Material and Methods

### Strains, media and culture conditions

The following fungal strains were used in this study: *C. albicans* strain SC5314 which is a human clinical isolate recovered from a patient with generalized candidiasis (32), several clinical isolates of *Candida* spp. received from the Fungus Testing Laboratory at the University of Texas Health Science Center at San Antonio: fluconazole-sensitive *C. albicans* CA1, CA4, and fluconazole-resistant *C. albicans* CA6, CA10, *C. glabrata* fluconazole-sensitive CG1, CG3 and fluconazole-resistant CG2*. C. parapsilosis* CP1, CP2, CP3, *C. neoformans* CN1, CN2, CN3, and *A. fumigatus* AF1, AF2, AF3. Some *Candida* strains were also obtained from the Division of Infectious Disease, Massachusetts General Hospital, Boston, MA: fluconazole-resistant strains of *C. albicans* CA2, *C. parapsilosis* CP4, CP5, *C. krusei* CK, and *C. tropicalis* CT2). The two *C. auris* isolates CAU-03 and CAU-09 were a kind gift from Dr. Shawn Lockhart, Centers of Disease Control (CDC), and the filamentous fungi including *Rhizopus delemar* 99.880 and *Rhizopus oryzae* 99.892*, L. corymbifera* 008049*, C. bertholletiae* 182, *M. circillenoides* 131 *and S. apiospermum* DI16-478 were a part of the fungal bank at Division of Infectious Diseases, Los Angeles Biomedical Research Center. All cultures were maintained by subculture on Yeast Peptone Dextrose media (YPD) at 37°C and stocks of these cultures stored in 20% glycerol at −80°C.

### HTS screening

Screening was performed at the Molecular Screening Shared Resource facility, at University of California, Los Angeles. A total of 50 μl of 1×10^4^ cells/ml fungal yeast cells (*C. albicans, C. auris*) or spores (*Aspergillus*) were suspended in RPMI-1640 supplemented with L-glutamine (Cellgro), buffered with 165 mM morpholinepropanesulfonic acid (MOPS), and plated into individual 384-well plates using an automated Multidrop 384 system (Thermo LabSystems). The New Prestwick Chemical Library consisting of 1233 drugs was used to pin one compound per well at 10 μM final concentration, using a Biomek FX liquid handler. Forty eight hours later the plates were scanned with a Flex Station II 384 well plate reader (Molecular Devices) to measure turbidity (OD600) of the wells. Molecules displaying >80% reduction in turbidity compared to control non-drug treated wells (MIC80) were considered as primary “hits”. Compounds commonly inhibiting all three fungal organisms were prioritized for planktonic dose response assays and for their activity against biofilm growth.

### Dose response assays

Dose response assay of AXD against planktonically grown fungi was determined in agreement with the CLSI M27-A3 (for yeast) and M38-A2 (for filamentous fungi) reference standards for antifungal susceptibility testing (13, 14). Each drug was used in the concentration range of 0.19 μg/ml to 24 μg/ml, and the MIC of AXD was compared to MIC of fluconazole, posaconazole or voriconazole, as controls. All strains described in Table 2 were tested at LA BioMedical Research Institute; however several of the *Candida* strains were also verified for their susceptibility to AXD independently at Massachusetts General Hospital. Inhibition of planktonic growth or filamentation due to drug treatment was also visualized and imaged using bright field microscopy. Microscopy was also used to directly visualize lateral yeast formation from planktonic *C. albicans* hyphae, or lateral yeast cells formed on the surface of the biofilms (dispersal) using microtiter plates.

### Biofilm growth and drug susceptibility testing

Biofilms of *Candida* spp., *C. neoformans*, and *A. fumigatus* were developed in 96-well microtiter plates, and susceptibility of the biofilm cells to AXD or thimerosal was carried out as described previously (33, 34). Biofilms were initiated either in the presence or absence of the drugs, or the drugs were tested on 48 h pre-formed biofilms, for efficacy evaluation. Inhibition of biofilm growth was measured by a standard calorimetric assay XTT that measures metabolic activity of the biofilm cells (18). Absorbance at 490 nm was measured using an automated plate reader. Biofilms formed by several other *Candida* spp. were further studied for their susceptibility to AXD.

Potential of AXD for synergistic use with fluconazole against *C. albicans* biofilms was investigated using a checkerboard assay, where dilutions of fluconazole (0.25 to 250 μg/ml) and AXD (0.3 to 2 μg/ml) were examined alone and in combination. Biofilm killing was measured by XTT assay. Drug concentration associated with 50% reduction in optical density compared to the no-drug control wells (EC_50_) was determined. The fractional inhibitory concentration (FIC) was then calculated as follows: [(EC_50_ of drug A in combination)/(EC_50_ of drug A alone)] + [(EC_50_ of drug B in combination)/(EC_50_ of drug B alone)]. Values of ≤0.5 revealed synergy, those of >0.5 but <2 indicated no interaction, and those of >2 were antagonistic (30).

### Mammalian cell toxicity assays

Primary human umbilical vascular endothelial cells (HUVEC) and human lung carcinoma derived A549 epithelial cell lines were used to determine the cytotoxicity of AXD. HUVEC cells were isolated and propagated by the method of Jaffe *et al*. (35). The cells were grown in M-199 (Gibco, Grand Island, N.Y) supplemented with 10% fetal bovine serum, 10% defined bovine calf serum and 2 nM L-glutamine, with penicillin and streptomycin. Second or third-passage endothelial cells were grown on collagen matrix on 96-well microtiter plates. Treatment with AXD was conducted in M-199 medium.

A549 cells were purchased from the American Type Culture Collection and grown in Dulbecco’s modified Eagle’s medium (DMEM) supplemented with 10% fetal bovine serum. A549 cells (1.5 × 10^5^/well) were used to seed 96-well plates and incubated at 37°C in a humidified atmosphere containing 5% CO_2_ for 24 h. The medium was then removed by aspiration, and the cells were washed twice with phosphate-buffered saline. Treatment with AXD was conducted with DMEM supplemented with 1% FBS.

Different concentrations of AXD in respected media were introduced into the cell lines, and incubated for 24 h at 37°C in 5% CO_2_. The extent of cellular damage to both cell lines, caused by AXD was quantified by a chromium release assay (36). Briefly, confluent mammalian cells were incubated overnight in respective media containing Na_2_^51^CrO_4_ (6 μCi per well; ICN Biomedicals, Irvine, Calif.). The next day, the unincorporated tracer was aspirated and the wells were rinsed three times with warm HBSS. Two hundred μl of media containing various concentrations of AXD (ranging from 0.12-59 μg/ml) was added to each well, and the plate was incubated for 24 h at 37°C in 5% CO_2_. At the end of the incubation, 100 μl of medium was gently aspirated from each well, after which the cells were lysed by the addition of 6 N NaOH. The lysed cells were aspirated, and the wells were rinsed twice with RadicWash (Atomic Products, Inc., Shirley, N.Y.). These rinses were added to the lysed cells, and the ^51^Cr activity of the medium and the cell lysates was determined. Control wells containing no drug were processed in parallel to measure the spontaneous release of ^51^Cr. After corrections were made for the differences in the incorporation of ^51^Cr in each well, the specific release of ^51^Cr was calculated by the following formula: (2X experimental release – 2X spontaneous release)/ (total incorporation – 2X spontaneous release).

### Cytotoxicity to immune cells

Wild-type C57Bl/6 primary bone marrow-derived macrophages were cultured by plating bone marrow cells in 50 ng/ml of M-CSF (Peprotech, Rocky Hill, NJ) in complete RPMI (RPMI 1640 with 2 mM L-glutamine, 10% heat-inactivated fetal bovine serum, and 1% penicillin-streptomycin) for 7 days, then counted and seeded at 1×10^5^ in 100 μl of complete PRMI overnight to allow for adhesion.

To examine cytotoxicity of AXD, bone marrow-derived macrophages were incubated in varying concentrations of AXD for 24h and stained with DAPI (Invitrogen, Carlsbad, CA) for viability assessment using an inverted epifluorescence microscope (Olympus IX70, Center Valley, PA) using 10X objective, with a X-cite 120 metal halide light source (EXFO, Mississauga, ON, Canada). Percent cell viability was determine using 1 – [DAPI-positive cells were divided by total cells by phase contrast] × 100.

AXD was also examined for its capacity to block proliferation of a human promyelocytic cell line, HL-60. Cells were stained with 2 mM CFSE (Carboxyfluorescein Succinimidyl ester) for 5 minutes and washed with 1X RPMI media three times. This dye is commonly used to measure cell proliferation; with each cell division the amount of CFSE is diluted in half, which can be observed via flow cytometry (37). After the staining, the cells were counted and adjusted at cell density of 5X10^6^ cells/ml and plated 100 μl/well in a round bottom 96-well plate. A two-fold serially diluted AXD was added in wells containing cells. The final drug concentration obtained was between 0.004 and 10 μg/ml. Drug-untreated and unstained cells in a number of wells were included as controls. Plates were incubated at 37°C for 48 hours to allow the cell proliferation. After 48 hours, the cells were collected and acquired in flow cytometer. The unstained cells were used to gate the CFSE positive HL60 cells. The shift in the peak of CFSE+ HL60 cells were considered proliferating cells.

### *In vivo* biofilm drug susceptibility

A mouse central venous catheter infection model was used for biofilm studies as previously described (31). These *in vivo* experiments were approved by the Los Angeles Biomedical Research Institute, Harbor-UCLA-IACUC. Briefly, we used catheterized 8-week old C57BL/6 male mice, purchased from Charles River labs (Wilmington, MA), where the surgery was performed. The surgery involves insertion of a Silastic catheter into the jugular vein of the mice. Patency is tested, and the catheter is filled with heparin lock solution and plug-sealed. Following receipt of the jugular vein catheterized mice, the catheters were instilled with 25 μl of *C. albicans* inoculum of 5×106 cells/ml (entire catheter volume) using a 23-gauge blunt-ended needle after removal of the plug and the lock solution (the plug will be put back in place after inoculation). Cells were allowed to develop biofilms for 24 h, after which the catheters were treated with 3 μg/ml AXD for 48 h. Biofilms growing in replicate mice catheters were also subjected to fluconazole (250 μg/ml) or caspofungin (0.125 μg/ml), as comparative controls. The catheters were cut laterally and imaged under a phase contrast microscope to visualize the morphology of the cells growing within the catheters of the individual groups. Additionally, the distal 2 cm of the catheters were into small pieces, vortexed vigorously and homogenized for plating on to YPD plates for viability count measurements.

### Statistical methods

All *in vitro* secondary assays were done in triplicate and repeated once. Experiments were conducted in a randomized fashion, and subjected to unpaired two-tailed t-tests and/or ANOVA with Kruskal-Wallis post-test to determine significance of results (for p ≤ 0.05). For *in vivo* studies, differences in catheter fungal burden between the four groups (6 mice per group), were presented as percent reduction in CFU in the individual drug treated groups compared to the control untreated mice. A two tailed t-test with a p-value of <0.05 was considered significant.

## Acknowledgment

We would like to thank Dr. Nathan Wiederhold, Director, Fungus Testing Laboratory, University of Texas Health Sciences Center at San Antonio, for providing us with several clinical isolates of fungi used in this study. These studies were funded by a grant from the National American Heart Association #16SDG30830012 to PU. ASI is supported by Public Health Service grant from the National Institutes of Allergy and Immunology R01 AI063503.

**S1:** Primary screening: MIC80 of alexidine dihydrochloride and thimerosal against pre-formed biofilms of three different fungi as determined by XTT reading (OD490). Both drugs kill biofilm cells at <10 μM concentration.

**S2:** Inhibitory effect of AXD against pre-formed biofilms of four different fluconazole resistant *Candida* spp. AXD kills preformed biofilms between 3-6 μg.

**S3A:** Chemical structure of Alexidine dihydrochloride

**S3B:** Inhibition of early cellular division of HL60 cells by AXD

